# Identification of neuronal lineages in the *Drosophila* peripheral nervous system with a novel multi-spectral lineage tracing system

**DOI:** 10.1101/618264

**Authors:** Macy W. Veling, Ye Li, Mike T. Veling, Christopher Litts, Nigel Michki, Hao Liu, Dawen Cai, Bing Ye

## Abstract

Elucidating cell lineages provides crucial understanding of development. Recently developed sequencing-based techniques enhance the scale of lineage tracing but eliminate the spatial information offered by conventional approaches. Multispectral labeling techniques, such as *Brainbow*, have the potential to identify lineage-related cells *in situ*. Here, we report *Lineage Tracker Bitbow*, a “digital” version of *Brainbow* that greatly expands the color diversity, and a suite of statistical methods for quantifying the lineage relationship of any two cells. Applying these tools to *Drosophila* peripheral nervous system, we determined lineage relationship between all neuronal pairs. Based on the refined lineage map, we explored whether distinct *cis*-regulatory elements are used in controlling the expression of a terminal selector gene in distinct lineage patterns. This study demonstrates *LT-Bitbow* as an efficient tool for *in-situ* lineage mapping and its potential in studying molecular mechanisms in the lineage context.

## INTRODUCTION

Cell lineage, which denotes the developmental history of a cell, is the conceptual framework of organism formation (Papaioannou, 2016; Stent, 1985). Modern genetic techniques have undergone evolutionary improvement, compared to early dye filling- or cell transplantation-based techniques for lineage tracing (Woodworth et al., 2017). For instance, sequencing-based methods can distinguish hundreds to thousands of uniquely barcoded lineages, though at the expense of the *in situ* spatial relationships of these lineages due to tissue disassembly (Raj et al., 2018; Schmidt et al., 2017; Spanjaard et al., 2018). In contrast, imaging-based methods, such as *Brainbow*, can preserve the spatial information *in situ* but their efficiencies are still limited by labeling diversities for unambiguous lineage tracing (Cai et al., 2013; Hadjieconomou et al., 2011; Livet et al., 2007; Pan et al., 2013). Hence, there is an urgent need to create novel labeling and detection methods that allow highly efficient lineage tracing while preserving spatial information.

*Brainbow*, a multi-spectral labeling technology, is designed to randomly express one of three or four fluorescent proteins (FPs) from a single cassette, thus creating stochastic labeling colors in neighboring cells or cell lineages (Cai et al., 2013; Hadjieconomou et al., 2011; Livet et al., 2007; Pan et al., 2013). When more color variants are needed, for example, to uniquely label many cell lineages, more than one *Brainbow* cassettes can be used to create differential expression levels of FPs. However, using color shades for lineage tracing is not always reliable. Distinguishing two color variants differing by subtle FP expression levels (e.g., color A generated from 2 YFP + 1 RFP + 1 CFP compares to color B generated from 1 YFP + 2 RFP + 1 CFP) can be challenging due to imaging noise (Cai et al., 2013). When applying *Brainbow* to trace cell lineages, daughter cells in the same lineage are assumed to inherit the same color generated in the mother stem cell. However, protein synthesis levels in daughter cells can be quite different. A more robust color generation mechanism is highly desirable for efficient and reliable lineage tracing.

One way to generate more robust *Brainbow* lineage labels is to localize the same FPs to different sub-cellular compartments. Cytoplasmic membrane-targeted and nucleus-targeted FPs, expressed through genome integration by electroporated transposase, have been used to differentiate neighboring neuronal lineages in chick and mouse embryonic brains (Garcia-Moreno et al., 2014; Loulier et al., 2014). However, transposase integrates various number of targeting cassettes in different cells, making it difficult to estimate the probability of each label combination for quantitative analysis. Generating transgenic animals with a fixed number of labeling cassettes would solve this problem. For instance, the *Raeppli* strategy utilized 4 FPs to create up to 4 × 4 = 16 membrane and nucleus color combinations in transgenic *Drosophila* (Kanca et al., 2014). Another recombination mechanism, implemented in the *CLoNe* and the *MultiColor FlpOut* (*MCFO*) *Drosophila* lines, generates random colors by stochastic removal of the expression stops from each FP module (Garcia-Moreno et al., 2014; Nern et al., 2015). For instance, a *MCFO* fly integrates 3 different stop-“spaghetti monster GFPs” (smGFPs) modules into 3 distinct genomic loci and generates up to 2^3^−1=7 smGFP combinations. However, further expanding the color pool requires inserting more FP modules to additional genomic loci to prevent inter-module recombination. In addition, the optimal color outcome can be difficult to obtain from *CLoNe* and *MCFO* animals. This is because while low FLP activity results in simple single-marker colors, high FLP activity often results in expressing all FPs in most of the cells (Nern et al., 2015). In conclusion, the small unique color pools generated by the above mentioned methods result in frequent observations that neighboring cells or lineages are labeled by the same color.

To overcome the limitations discussed above, we introduced a novel “digital” format of *Brainbow* for robust lineage tracing, which we termed *Lineage Tracker Bitbow* (*LT-Bitbow*). A single *LT-Bitbow* cassette composites binary switches to independently determine an ON/OFF state of 5 FPs to generate up to 2^5^-1=31 color variants. We applied *LT-Bitbow* to trace neuronal lineages in the *Drosophila* peripheral nervous system (PNS). We chose the fly PNS as a model because (1) individual PNS neurons are identifiable by their soma locations and neurite patterns; (2) PNS neuronal lineages reported in previous studies can serve as references for result comparison (Brewster and Bodmer, 1995, 1996); and (3) previous neuronal lineage mapping in PNS was incomplete due to technical challenges. As there is a lack of statistical methods for color-based lineage tracing, we developed a lineage relatedness test to determine the likelihood of two cells being lineage-related based on their colors. Using this statistic test, we confirmed and rejected previously determined lineages, as well as revealed unknown lineages. Moreover, applying *LT-Bitbow* labeling at different developmental time points, we revealed the proximate birth-timing of many PNS neuronal lineages. Based on the refined lineage map, we had the opportunity to examine whether distinct *cis*-regulatory elements are used in controlling the selective expression of *dendritic arborization 1* (*dar1*), a terminal selector gene for multipolar neurons derived through distinct lineage patterns (Cajal, 1995; Wang et al., 2015).

## RESULTS

### Lineage Tracker-Bitbow

As opposed to previous *Brainbow* designs that expand color variations by recombining multiple cassettes to mix different intensity levels in each spectral channel, *Bitbow*, a “digital” format of *Brainbow*, utilize several (“N”) FPs, and randomly assign each to an ON or OFF expression state upon FLP-mediated recombination (Figure 1A). As a result, *Bitbow* generates “N-bit” (2^N^-1, minus the all-OFF state) “colors” from a single cassette. We further optimized *Bitbow* for lineage tracing and generated a *Lineage Tracker-Bitbow* (*LT-Bitbow*) cassette containing 5 FPs (mAmetrine, mTFP, mNeonGreen, mKusabira-Orange2 and tdKatushka2). Individual FP was fused to a human histone H2B (hH2B) for nuclear tagging. The cDNA of each nuclear-FP was positioned in the inverse direction and flanked by a pair of incompatible FRT sites, thus permitting spinning the FPs in forward or inverse orientation upon FLP recombination (Cai et al., 2013; Mcleod et al., 1986; Schlake and Bode, 1994; Turan et al., 2010; Volkert and Broach, 1986). Unlike previous *Brainbow* designs, recombination of each nuclear-FP in *LT-Bitbow* was independent from each other, resulting in up to 2^5^-1=31 unique colors by a single cassette (Figure 1B). Finally, a UAS sequence and a p10 baculovirus poly-adenylation sequence were placed upstream and downstream, respectively, to each of the five FP recombination units to allow strong FP expression when driven by a Gal4 driver (Pfeiffer et al., 2012). We chose the FPs whose native signals are bright and stable so that immunostaining is not required. In addition, spectral imaging and linear unmixing were applied and optimized for separating the five fluorescent signals to avoid signal bleed-through (Figure 1C).

**Figure 1.**
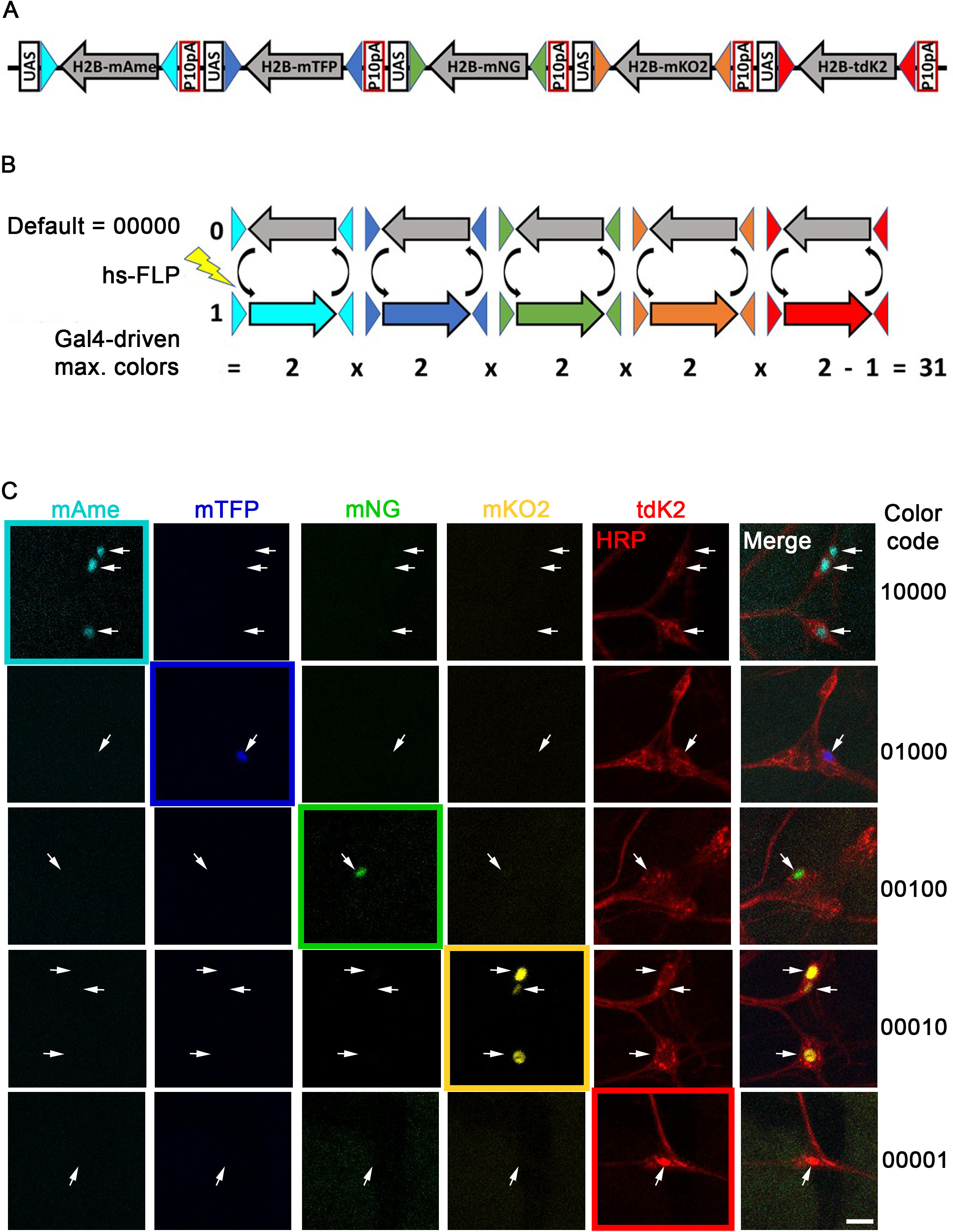
Design and color-coding principles of *LT-Bitbow*. A. Schematic of the *LT-Bitbow* “spinning” design. The H2B fusion nuclear-FP (nucFP) open reading frames are positioned in the reverse direction as a default non-detectable color in the absence of FLP.
B. Schematics of color-coding in *LT-Bitbow*. Upon heat-shock-induced FLP expression, which causes recombination of FRT sites, each H2B-FP module can stochastically and independently spin into a forward (1, visible) or reversed (0, non-detectable) expression state. Maximum of 31 visible colors can be detected in the presence of a Gal4 driver.
C. Spectral imaging and linear unmixing are performed to identify the five-bit *LT-Bitbow* color code for each cell, in which “1”s represent the FPs that are turned on and “0”s represent those that are not turned on. HRP: anti-HRP immunostaining, which specifically labels PNS neurons. Scale bar: 10 μm.

To evaluate the labeling efficiency of *LT-Bitbow* in the *Drosophila* larval PNS, we first combined a transgene encoding heat-shock-inducible flippase (*hsFLP*) and the pan-neuronal driver *elav-Gal4* to determine how often neurons expressed colors in the 3^rd^ instar larvae. Without any heat-shock, an average of 2.89 ± 0.90% (mean ± standard error of mean) of PNS neurons showed colors when the larvae were raised at 23 ℃ (Figure 2A), likely due to leaky FLP expression (Golic and Lindquist, 1989). Lowering the developmental temperature to 18 °C did not reduce this leaky FLP activity (data not shown). Induction of FLP expression by a 30 min heat-shock at 37 °C that started at the 3^rd^, 4^th^, or 5^th^ hours (hr) after egg laying (AEL) led to significant increases of the number of PNS neurons that expressed colors (up to 18.62 ± 2.74 %) (Figure 2A). Heat-shock that started at 2^nd^ hr AEL turned on color expression in similar number of neurons compared to the non-heat-shock condition (Figure 2A), possibly due to either low levels of FLP expression or poor accessibility of FLP to the *LT-Bitbow* construct at very early stages of embryogenesis. Interestingly, we observed that the efficiency of turning on colors at the 6^th^ hr AEL decreased to the level of non-heat-shock condition. This may indicate a lower efficiency of FLP recombination in the post-mitotic cells (detailed below).

**Figure 2.**
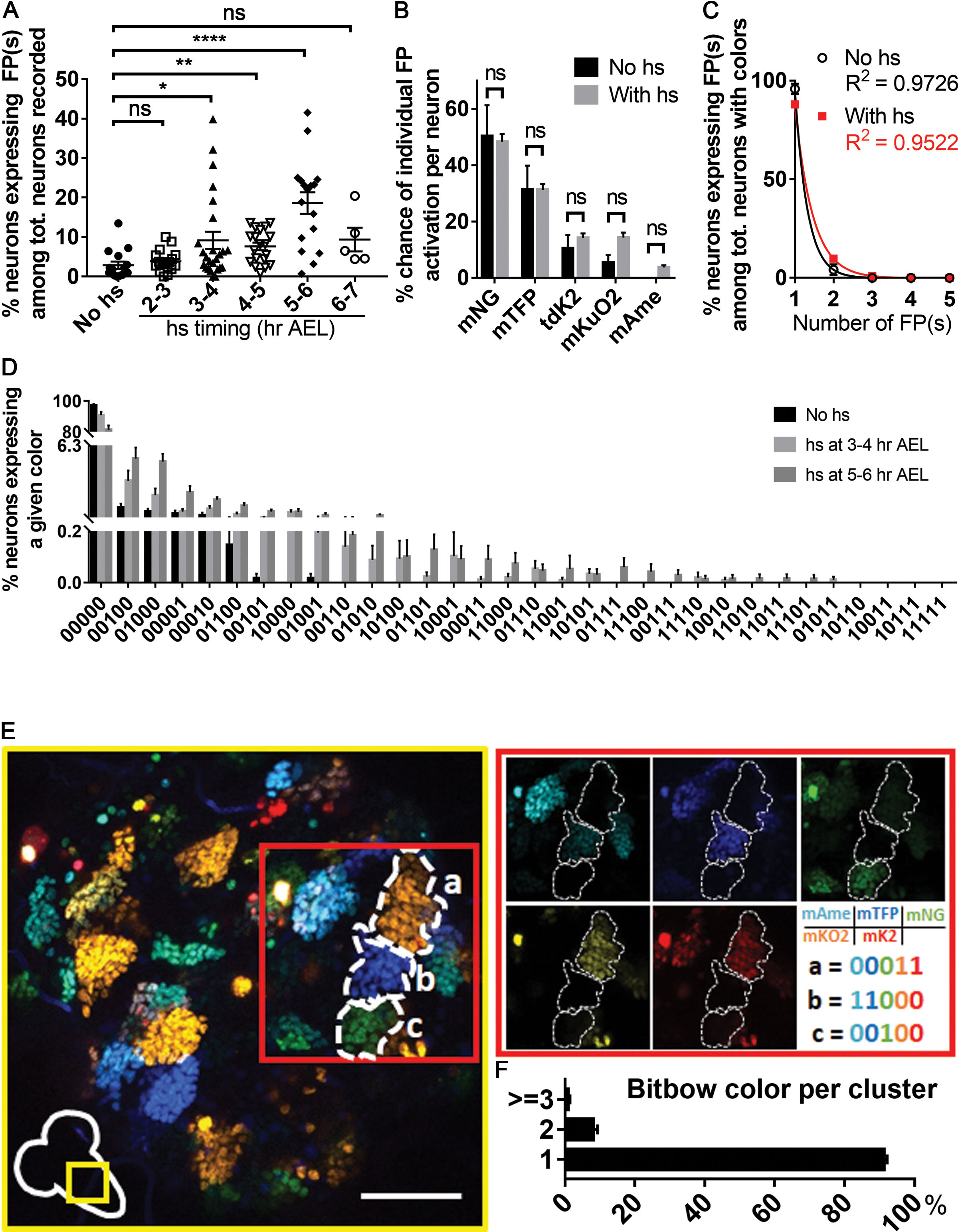
Labeling and lineage tracing efficacy of *LT-Bitbow* in *Drosophila* nervous system. A. Quantification of the labeling coverage of *LT-Bitbow* by heat-shock FLP induction in the PNS at various time points during early embryonic development. Sample numbers (larvae): no hs, 16; 2-3 hr AEL, 18; 3-4 hr AEL, 25; 4-5 hr AEL, 19; 5-6 hr AEL, 17; 6-7 hr AEL, 5. P-values were calculated by Dunn’s multiple comparisons test. ns: p > 0.05, *: 0.01 < p ≤ 0.05, **: 0.001 < p ≤ 0.01, ***: 0.001 < p ≤ 0.001, and ****: p < 0.0001.
B. Comparison of the turn-on efficiency of each FRT-nucFP-FRT module in the PNS. Sample numbers (larvae): no hs, 16; hs, 81. P-values were calculated by multiple comparisons using the Holm-Sidak method.
C. The percentage of neurons that express more than one nucFP species decreases exponentially in the PNS. This indicates that the recombination of each FRT-nucFP-FRT module is independent to each other. Sample numbers (larvae): no hs, 15; hs, 81. P-values were calculated by multiple comparisons using the Holm-Sidak method.
D. 27 color codes were observed with different frequencies in the PNS from a *Drosophila* line that carries only one *LT-Bitbow* cassette. Sample numbers (larvae): no hs, 15; 3-4 hr AEL, 25; 5-6 hr AEL, 17.
E. Representative images of *LT-Bitbow* labeling in thoracic segments in the CNS of 3^rd^ instar larvae with embryonic heat-shock FLP induction. Most of the cell clusters (i.e. lineage clones) were labeled by the same color. 193 clusters were analyzed. Scale bar: 50 μm.
F. Proportion of thoracic lineage clones labeled by one, two or more *Bitbow* color tags, quantified across n=193 clusters. P-values were calculated using unpaired two-tailed t-test and α of 0.05 is used as the cutoff for significance. Error bars: standard error of mean.

As different FRT sites exhibit different efficiencies of FLP-induced recombination (Schlake and Bode, 1994; Turan et al., 2010), we quantified the expression frequency of individual FPs. mNeonGreen displayed the highest expression frequency, followed by mTFP, mKusabira-Orange2, tdKatushka2, and mAmetrine (Figure 2B). Next, we quantified the frequency of neurons expressing one, two, three, four, or all five FPs. We found that such frequency decreased exponentially from expressing one FP to five FPs, which suggests that the recombination of distinct FRT sites is independent of each other as the expression of one FP does not appear to increase the chance of expressing other FPs (Figure 2C). Finally, the expression frequencies of all the FP combinations (i.e., color variants) are quantified (Figure 2D) for our lineage relatedness test (detailed below). Despite that different combinations exhibited different frequencies, we observed 27 of the 31 expected colors after heat-shock-induced FLP expression.

One potential drawback of *LT-Bitbow* for lineage tracing is that the “spinning” recombination design may cause color changes over time if the leaky FLP activity is persistent in a cell. As a result, cells with altered colors in the same lineage may be mis-identified as in distinct lineages. To evaluate whether *LT-Bitbow* is reliable for lineage tracing, we performed the same heat-shock experiment and examined the color labeling in the central nervous system (CNS). Unlike the sensory organ precursors (SOPs) in the PNS, neuroblasts (NBs) in the CNS divide many more times to generate cell clusters with clear boundaries that can be easily separated from each other even with simple GFP labeling (Birkholz et al., 2015; Truman and Bate, 1988). We found that the vast majority of the cells in each cluster were labeled in the same *LT-Bitbow* color (Figure 2E). We occasionally observed a few scattered cells labeled by different colors in a cluster, and rarely observed a cell cluster that was labeled by two distinctly colored clones (Figure 2F). This result indicates that the embryonic heat-shock induced *LT-Bitbow* labeling mostly happens in neural stem/progenitor cells and that color changes caused by persistent FLP activity is minimal. Indeed, we also observed that *LT-Bitbow* had the lowest labeling efficiency in the PNS when heat-shocked at the 6^th^ hr AEL (Figure 2C), a stage that has more post-mitotic cells. Taken together, these results suggest that *LT-Bitbow* provides more unique colors from a single cassette than previous variants of *Brainbow* and is reliable for lineage tracing in the *Drosophila* nervous system.

### Identification of neuronal lineages in the *Drosophila* larval PNS by *LT-Bitbow*

*Drosophila* PNS neurons are generated through lineages that repeat in the abdominal hemi-segments (Figure 3A) (Hartenstein and Campos-Ortega, 1984; Hartenstein et al., 1987). The SOPs in each hemi-segment are born and delaminated from ectoderm at approximately 3.5 to 7.5 hr AEL to eventually give rise to 45 PNS neurons (Bodmer et al., 1989; Ghysen and Okane, 1989; Younossi-Hartenstein and Hartenstein, 1997). We therefore induced *LT-Bitbow* recombination in PNS neurons by heat-shock at 3-4 or 5-6 hr AEL and recorded their colors in 3^rd^ instar larvae (Figure 3B). We expect to observe a significantly higher probability of two neurons being labeled in the same colors if they belong to the same lineage (i.e., inheriting the same colors from their SOP, Figure 3C) than if they belong to different lineages (i.e., acquiring the same colors by chance). Based on the Cohen’s Kappa (κ) statistics, we designed a randomized lineage relatedness test to determine whether a pair of two neurons are lineage-related (Cohen, 1960; Dwass, 1957; Nichols and Holmes, 2007; Nichols and Holmes, 2002). Ranging from −1 to 1, κ = 1 indicates the two neurons are always labeled by the same colors, κ = 0 indicates the neurons are labeled by the same colors by chance, and κ = −1 indicates the two neurons are always labeled by different colors. We calculated the κ value of every pair of PNS neurons, and determined whether each κ value is significantly different against the probability of labeling the same neuron pair with the same colors by chance in a randomization test (Figure 3D, and see STAR methods for further details). Finally, a ≤ 2% false discovery rate (FDR) based on the Benjamini Hochberg method was applied to confirm any pair of neurons that are lineage-related (Benjamini and Hochberg, 1995).

**Figure 3.**
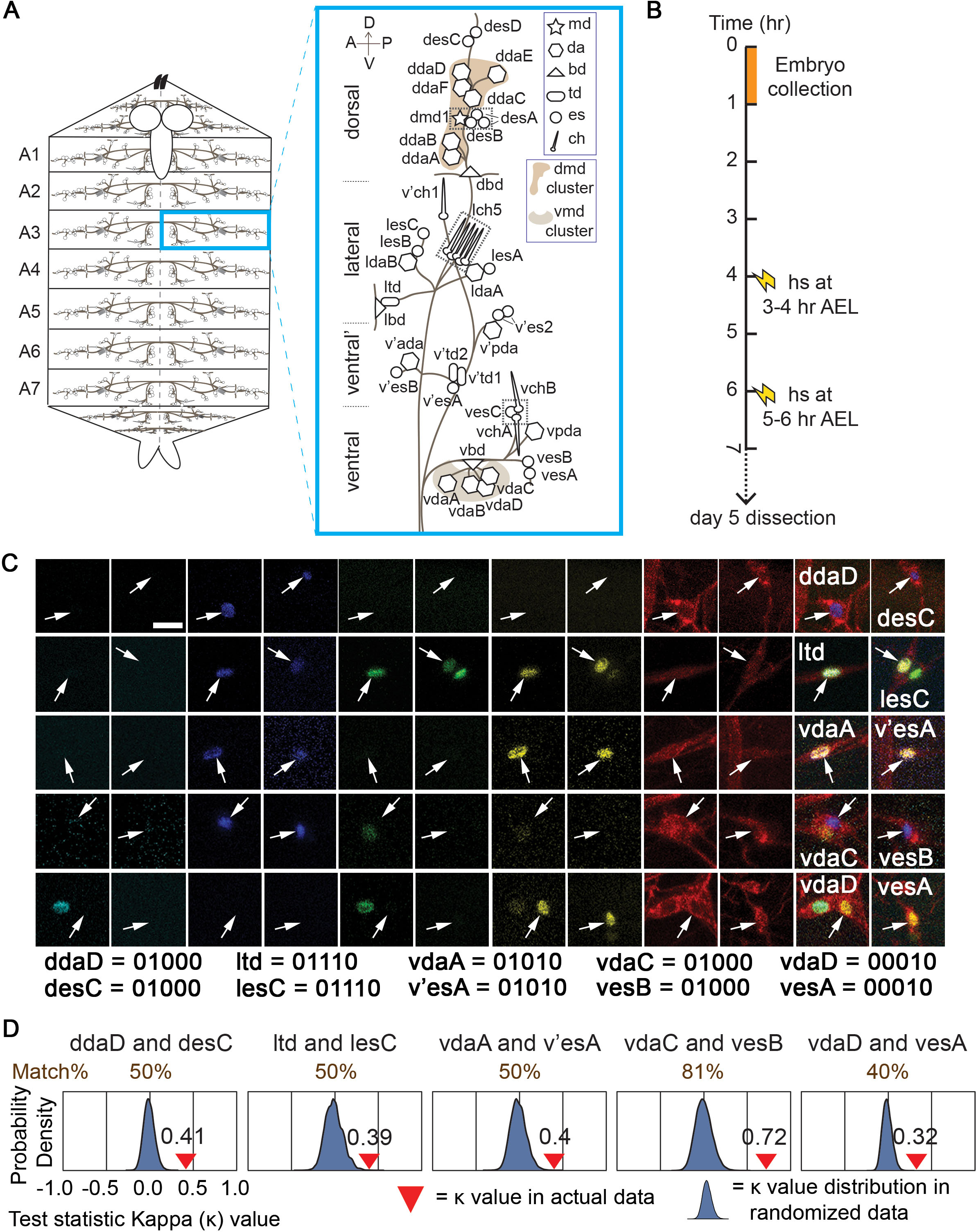
*LT-Bitbow* efficiently identifies neuronal lineages of *Drosophila* PNS neurons. A. Schematic drawing of the distribution of PNS neurons in one hemi-segment of the larval body wall. In the PNS, multipolar neurons include the md, da, bipolar dendrite (bd), and td types, whereas bipolar neurons include the es and chordotonal (ch) types.
B. Schematic drawing of the experimental steps.
C. Representative fluorescent images that show matching color-codes of pairs of neurons previously thought to be independent to each other. These newly identified neuronal pairs are the five es neurons (desC, lesC, v’esA, vesA and vesB) and their lineage-related da or td neurons. These es neurons have been previously reported as the only neuron from individual SOPs (Brewster and Bodmer, 1995). Larvae were heat-shocked at 3-4 hr AEL. Scale bar: 10 μm.
D. Statistical analysis confirms that these neuron pairs are lineage-related. The κ values calculated based on experimental observations (red arrows) are significantly higher (i.e., closer to “1”) than those calculated based on a random color distribution (blue). (See also Figure S1 and S2.)

Since some neurons are labeled at the 3^rd^ instar larval stage without heat-shock, we first examined how this leaky FLP activity would affect the outcomes of our color-based lineage analysis. From the non-heat-shock experiments, our lineage relatedness test determined that all the neurons appear to be in distinct lineages (Figure 4, left panel). This indicates that the leaky FLP activity is unlikely to recombine the *LT-Bitbow* cassette in any SOP until accumulating to a sufficiently high level to recombine in the post-mitotic neurons. This also suggests that our lineage relatedness test is reliable to avoid the false-positive assignment of two cells being lineage related.

**Figure 4.**
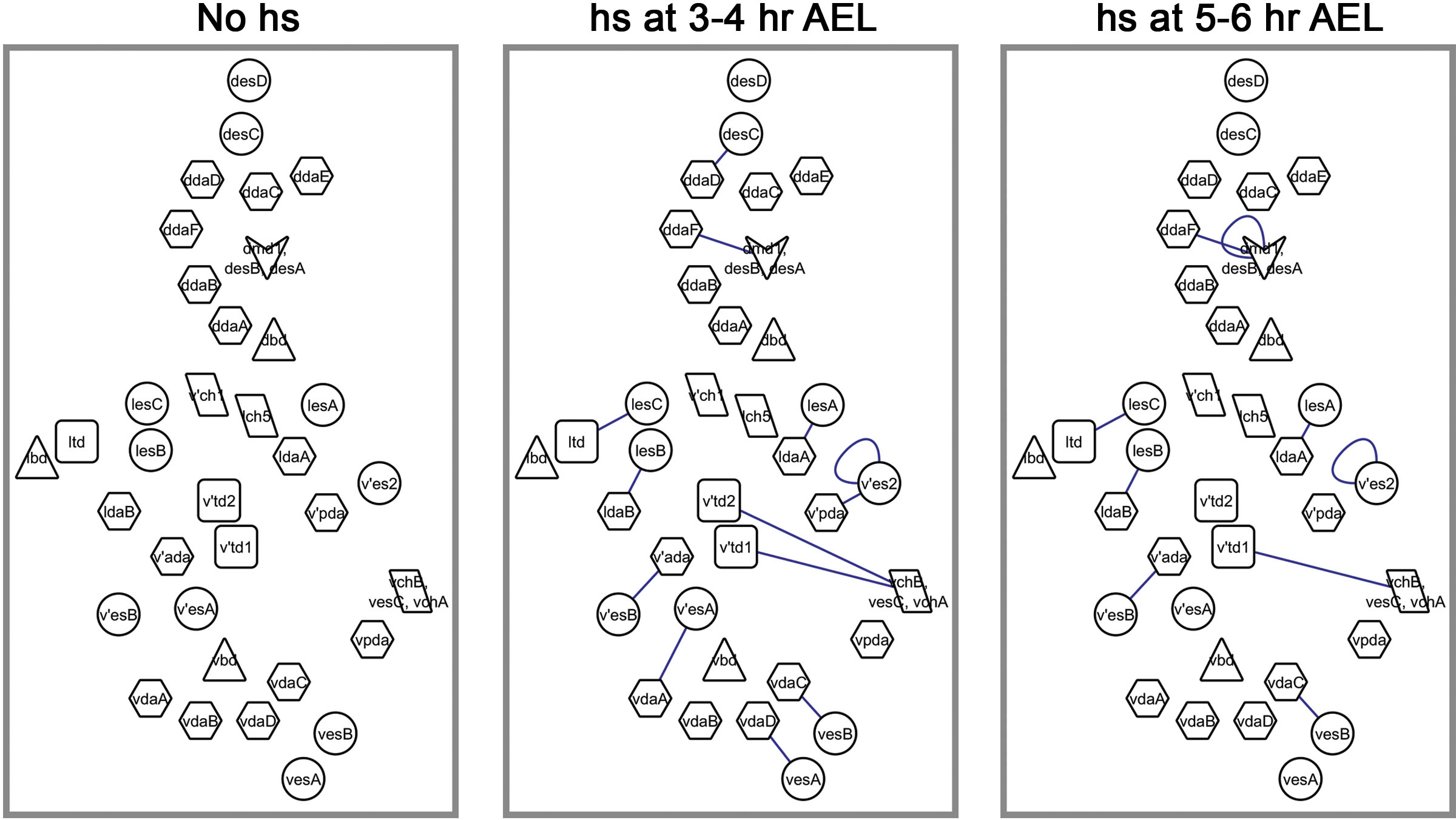
PNS lineage maps revealed by *LT-Bitbow* at different developmental times. Shown are neurons in a hemi-segment. These maps are based on the lineage relationships of all neurons determined by their calculated κ values from experiments with no heat-shock (left), heat-shock at 3-4 hr AEL (middle), and 5-6 hr AEL (right). Each line connects a pair of lineage-related neurons with ≤ 2% false discovery rate (FDR). Three groups of neurons that cannot be identified individually (dotted boxes in Figure 3A) are presented as single symbols in these maps: the arrow head includes four neurons—dmd1, desB, and 2 desA—which are located in close proximity; the parallelogram labeled as “lch5” includes 5 ch neurons; and the parallelogram labeled as “vchA, vchB, vesC” includes the three neurons indicated. (See also Figure S3.)

Previous studies used heat-shock to sparsely induce lacZ expression at early developmental time points and referred cell clusters that are located nearby to be in the same SOP lineages (Brewster and Bodmer, 1995, 1996). We started *LT-Bitbow* lineage tracing with heat-shock at 3-4 hr AEL. The lineage relatedness test led to some conclusions the same as the previous studies did: 4 distinct lineages give rise to a pair of ldaA-lesA, ldaB-lesB, v’ada-v’esB, or v’pda-v’es2 neurons, respectively (Figure S1).

The lacZ-based lineage tracing approach may incorrectly determine neurons in different lineages as in the same lineage, if they are easily labeled at the same heat-shock time point. *LT-Bitbow*, on the other hand, labels the lineage-unrelated neurons in distinct colors even if their progenitors are labeled at the same heat-shock time point. For instance, prior studies suggested that some of the dendritic arborization (da) neurons in the dorsal multidendritic (dmd) cluster were generated from one SOP and those in the ventral multidendritic (vmd) cluster were generated from another SOP (Figure 3A, shaded clusters) (Brewster and Bodmer, 1995, 1996). In contrast, our results showed that none of the da neurons from either dmd or vmd cluster are lineage related (Figure S2).

The lacZ-based lineage tracing approach may also incorrectly determine neurons in the same lineage as in different lineages, if they migrate away from each other. *LT-Bitbow*, on the other hand, relies on color identify but not spatial proximity that avoids this type of error. For instance, the 5 external sensory (es) neurons (desC, lesC, v’esA, vesA and vesB) had been previously concluded as the only neurons generated from 5 SOPs. Our lineage relatedness test revealed that each of the 5 es neurons were lineage-related to either a da or a tracheal dendrite (td) neuron (ddaD-desC, ltd-lesC, vdaA-v’esA, vdaC-vesB, and vdaD-vesA, Figure 3C and 3D). This finding is in agreement with the results mentioned above that dda or vda neurons in the dmd or vmd clusters, respectively, are not lineage-related to each other, but are lineage-related to the des or ves neurons, respectively (Figure 3C, 3D and Figure S2).

### Estimating the birth-timing of neurons by *LT-Bitbow*

One inherent benefit of using *LT-Bitbow* for lineage mapping is its efficiency. *LT-Bitbow* does not require very sparse labeling. Indeed, all samples expressing any FP colors were utilized in the analysis. This allowed us to efficiently map the complete PNS lineages using only 25 larvae for each heat-shock time point.

Comparing experiments done with 3-4 hr AEL heat-shock to those done with 5-6 hr AEL heat-shock, we found that even though a lot more neurons were labeled (Figure 2A), the overall percentage of neuron pairs that were labeled in matching colors decreased from 28.80% to 23.98%, respectively. Consistently, the lineage relatedness tests indicated that the ddaD-desC, v’pda-v’es2, v’td2-vchA/vchB/vesC, v’esA-vdaA, and vdaD-vesA neuron pairs no longer appeared to be lineage-related when heat-shock was applied at 5-6 hr AEL (Figure 4, middle and right panels). Interestingly, while the frequency of color matching of the v’ada-v’esB pair or the vdaC-vesB pair dropped significantly in the experiments done with 5-6 hr AEL heat-shock, that of the ldaA-lesA pair or the two es neurons in the v’es2 group remained unchanged (Figure S3). These observations suggest that at least one of the neurons in the v’ada-v’esB or the vdaC-vesB lineage finish its final cell division earlier than those in the ldaA-lesA or the v’es2 lineage.

Taken together, the birth-timing of individual lineages can be estimated by comparing the color matching frequency between earlier and later heat-shock time points.

### Different combinations of *cis*-regulatory elements determine the expression of the terminal selector gene *dar1* in different lineages

Neurons assume distinct morphologies that can be categorized into multipolar, bipolar, and unipolar subtypes (Cajal, 1995; Wang et al., 2015). We previously found that *dar1* is selectively expressed in post-mitotic multipolar neurons and is necessary and sufficient for the multipolar morphology (Wang et al., 2015), thus functioning as a terminal selector for multipolar neuronal morphology (Hobert, 2008). With a refined PNS neuron lineage map, we can now investigate whether distinct *cis*-regulatory elements control the expression of *dar1* in multipolar neurons that are generated through distinct lineage patterns (e.g., da/es, da/es/es, or solo-da) (Figure 4).

We first created GFP-reporter *Drosophila* lines to identify the minimal genomic region that specifies *dar1* expression in multipolar neurons in the PNS. We found that an approximately 500 base-pair (bp) region located about 950 bp upstream from the *dar1* transcription start site was sufficient for driving GFP expression in most multipolar neurons, in particular nearly all da neurons except vdaD (Figure 5A-B and S4A). Interestingly, this 500-bp fragment also drives GFP expression in all bipolar ch neurons, but rarely in bipolar es neurons (Figure 5B). Moreover, consistent with the fact that *dar1* is absent in the larval CNS, which only contains non-multipolar neurons, this 500-bp fragment only drove GFP expression in a few neurons in the entire CNS, which was similar to a control driver that did not contain any *dar1* fragment (Figure S4B). These observations collectively suggest that this 500-bp fragment drives *dar1* expression in multipolar neurons while additional regulatory mechanisms prevent *dar1* expression in ch neurons.

**Figure 5.**
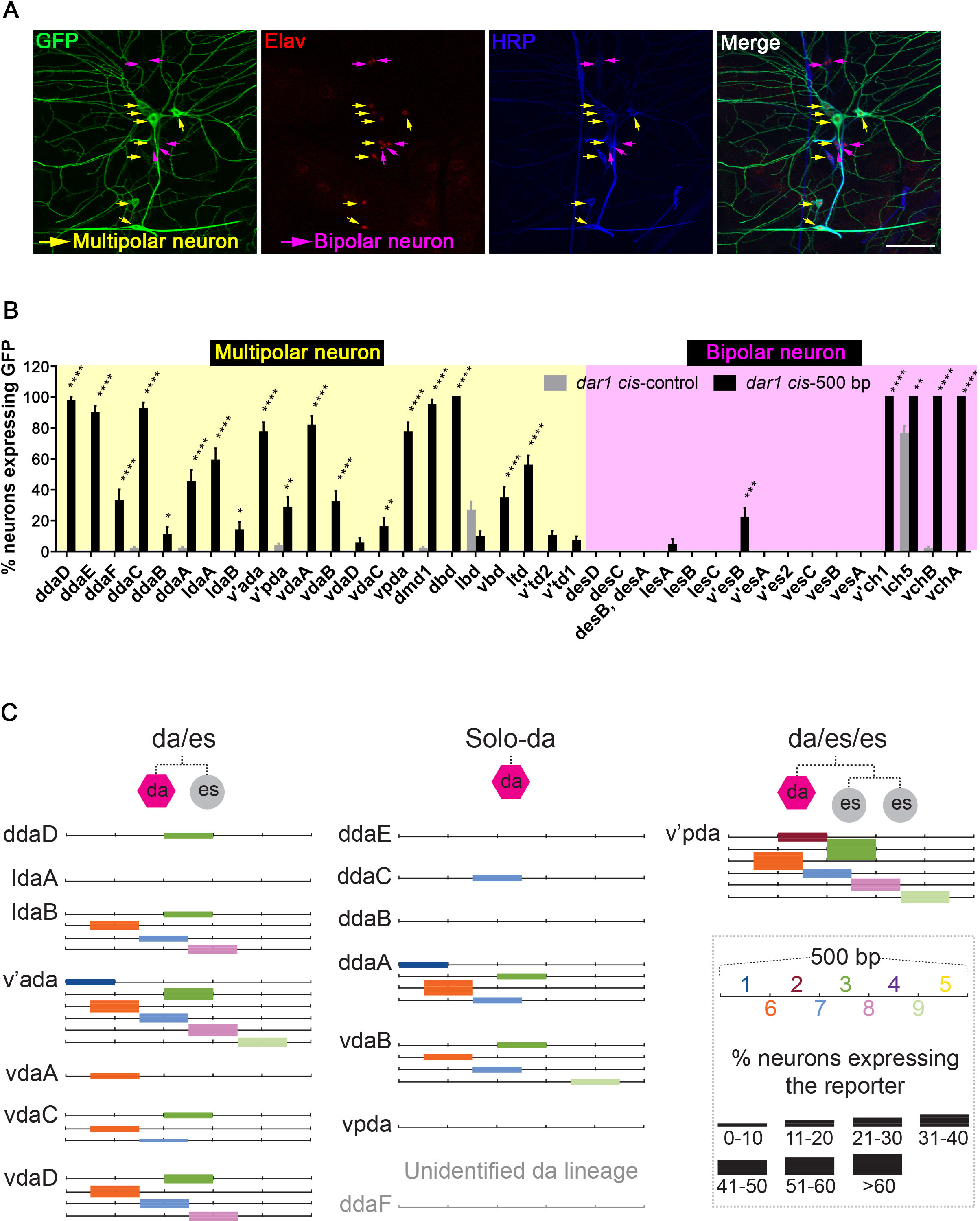
Multipolar neurons arising from different lineages express the morphological terminal selector gene *dar1* through different combinations of *cis*-regulatory elements. A. A 500-bp genomic DNA upstream of *dar1* transcription initiation site drives the expression of GFP-reporter in multipolar neurons (yellow arrows) but not in bipolar neurons (magenta arrows). Green: membrane-bound GFP reporter; red: pan-neuronal marker Elav; blue: anti-HRP. Scale bar: 50 μm.
B. Quantification of the percentage of PNS neurons expressing GFP in the 500-bp driver line (“*dar1* cis-500 bp”). PNS neurons (lbd and lch5) are excluded in the following studies due to high non-specific reporter expression in the control transgenic animals (“*dar1* cis-control”). P-values were calculated by multiple comparisons using the Holm-Sidak method. *: 0.01 < p ≤ 0.05, **: 0.001 < p ≤ 0.01, ***: 0.001 < p ≤ 0.001, and ****: p < 0.0001. Error bars: standard error of mean.
C. da neurons are categorized by three lineage patterns: da/es, da/es/es, and solo-da. The black lines represent the 500-bp genomic DNA fragment that drives reporter expression. Individual color block represents the ∼100 bp fragments that drive reporter expression significantly in subsets of neurons. The thickness of the color blocks approximates the percentage of neurons expressing the reporter. (See also Figure S4.)

Next, we set to determine whether this 500-bp fragment contains *cis*-regulatory elements that specifies *dar1* expression in certain lineage patterns of da neurons. To do so, we synthesized 9 overlapping DNA fragments of ∼100 bp each to tile the entire 500-bp (Figure S4A). We then generated 9 reporter *Drosophila* lines, in each of which GFP was driven by one of the ∼100 bp fragments. We then quantified the GFP expression pattern in da neurons (Figure 5C). We found that except fragments 4 and 5, all the other ∼100-bp fragments drove GFP expression in at least one da neuron, indicating that they contain at least one *cis*-regulatory element. We also found that none of the 100-bp *cis*-regulatory fragments drove GFP expression in any ldaA, ddaB, ddaE, ddaF or vpda neurons, which suggests that *dar1* expression in these neurons requires combinations of these *cis*-regulatory elements. Next, we examined whether any particular combination of *cis*-regulatory elements drives GFP expression in da neurons generated with the same lineage pattern. We found that a region tiled by fragments 3, 6, 7 and 8 drove common GFP expression in ldaB and vdaD, both of which are generated with the da/es lineage pattern. None of the other combinations of *cis*-regulatory elements drove a common GFP expression pattern between any two da neurons.

These results suggest that da neurons generated through the same lineage pattern do not necessarily achieve their morphologies through a common set of *cis*-regulatory elements. Instead, different combinations of *cis*-regulatory elements determine the basic morphological types in different lineages.

## DISCUSSION

Our findings demonstrate that the novel *LT-Bitbow* and the lineage relatedness test are proper tools for mapping neuronal lineages and estimating the birth-timing of neurons in the *Drosophila* nervous system. Providing limited unique color choices in a transgenic animal, previous multi-spectral labeling methods were constrained to study lineages that contain large number of cells forming physically connected clones (Loulier et al., 2014). *LT-Bitbow* overcomes this limitation by generating ∼30 reliably detectable FP combination colors. The different frequency of expressing each color becomes an advantage for lineage tracing because our lineage relatedness test weights in the rarely generated color in a modified κ statistics to reveal the lineage relationship between two neurons.

Mapping lineages by *LT-Bitbow* is much more efficient compared to previous method. For example, a previous study examined over 5000 embryos to identify the lineage relationships in the *Drosophila* larval PNS. This is because in order to reliably count the frequency of neighboring cells expressing a lacZ marker that was induced by heat-shock in a single progenitor cell, labeling needs to be extremely sparse (Brewster and Bodmer, 1995, 1996). Using *LT-Bitbow*, we achieved a similar goal with 40 animals (23 for heatshock at 3-4 hr AEL and 17 for heatshock at 5-6 hr AEL). In addition, we took advantage of the fact that each larval PNS neuron is identifiable by its cell body position and the morphology of proximal dendrites (Bodmer and Jan, 1987; Grueber et al., 2002; Jan and Jan, 1982), and combined neuronal morphology staining with *LT-Bitbow*’s multi-color nuclear labeling to assign each identified neuron to a lineage.

*LT-Bitbow*’s ability to generate many more unique colors also overcomes a fundamental limitation of the previous methods. With limited color choices, previous monochromic or multicolor labeling methods rely on physical proximity when determining whether two or more cells belong to the same lineage. This may generate false negative or false positive results; for instance, cells that migrate far from their sisters may be missed as in distinct lineages, or cells that locate close to it cluster boundary may be included in the neighboring lineages. Our method minimizes these errors by providing reliable statistics to determine whether two cells are lineage-related based solely on *LT-Bitbow* color but not on their physical locations. For instance, we discovered that even though da neurons in the vmd or dmd clusters are physically very close to each other, they are all generated by distinct progenitor cells (Figure S2). Interestingly, each of these da neurons was found to be lineage-related to a non-multipolar neuron that locates further away (Figure 3C-D).

Initiating *LT-Bitbow* labeling at different developmental time points permitted convenient birth-timing analysis in the context of lineage-relationship among PNS neurons (Figure 4). We found that several PNS neuronal pairs that were identified as lineage-related in experiments with an earlier heat-shock time point were identified as lineage-unrelated in experiments with a later heat-shock time point. This suggests that none of the neurons in those lineages were born at the earlier time point so that they inherited the same colors from their mother SOPs when heat-chock was applied. On the other hand, one of the neurons in any of these lineages became to complete its last cell division at the later time point so that its color is distinct from its sister neuron. However, this method could not be used to determine the birth-timing of those are the sole neurons in their lineages. Since the *LT-Bitbow* expresses bright nuclear targeted FPs, it is possible that the birth-timing of these neurons can be revealed by time lapse imaging in living larvae.

The success of our PNS neuronal lineage mapping created new opportunities for studying other biological aspects in the context of lineage development. For instance, it lays the foundation for studying whether particular molecular mechanisms are specified in neurons arising from different lineage patterns. Our results suggest that the transcriptional control of *dar1* gene in different da neuron lineages is complex. We speculate that distinct transcription factors are recruited to control *dar1* expression in different da neuron lineages, but common factors might also be used in certain subsets of da neurons. Profiling the expression pattern of transcription factors in the context of da neuron lineages may reveal the regulatory mechanisms underlying neuronal morphogenesis in different lineages.

In summary, we expect these new tools to greatly increase the efficiency and accuracy for studying *Drosophila* lineage development beyond the nervous system. These new tools also bring the potentials of combining lineage mapping with other fields of research to answer complex questions in developmental biology.

## ACKNOWLEDGEMENT

We thank Erica Edwards, Marya Ghazzi, Yimeng Zhao, and Tiffany Chen for aiding the creation of *LT-Bitbow* fly lines, Cheng-yu Lee, Scott Barolo, Catherine Collins for discussions, and Carl Zeiss Microscopy, LLC for microscopy support. This work was supported by the National Institutes of Health (R21MH106151 to D.C. and B.Y., R01MH110932 and R01AI130303 to D.C., R01MH112669 to B.Y., F31NS100391 and T32GM007315 to M.W.V.).

## AUTHOR CONTRIBUTIONS

M.W.V., B.Y. and D.C. conceived the project. B.Y. and D.C supervised the project. M.W.V., Y.L., C.L., M.T.V., and H.L. performed the experiments. Y.L. and D.C. designed and generated *LT-Bitbow Drosophila* lines. M.W.V., Y.L., D.C. and B.Y. designed the experiments of lineage tracing in larval PNS. M.W.V. and B.Y. designed the experiments for *dar1* expression regulation. M.W.V., W.T.V., N.M., and D.C. designed the statistical methods, and carried out the analysis. M.W.V, M.T.V., D.C., and B.Y. wrote the paper.

## STAR METHODS

### The *LT-Bitbow* construct and transgenic *Drosophila* lines

cDNAs encoding the following FPs were used: mAmetrine, mTFP, mNeonGreen, mKusabira-Orange2, and tdKatushka2 (Ai et al., 2008; Ai et al., 2006; Sakaue-Sawano et al., 2008; Shaner et al., 2013; Shcherbo et al., 2009). Human histone 2B protein (hH2B) was fused in frame to the N-terminus of individual FPs (hH2B-FP). Individual incompatible FRT sequence pairs (FRT^F3^, FRT^F14^, FRT^545^, FRT^F13^, or FRT^5T2^) was then placed in the opposing direction on both ends of the hH2B-FP sequence (Cai et al., 2013; Mcleod et al., 1986; Schlake and Bode, 1994; Turan et al., 2010; Volkert and Broach, 1986). And then each of the five FRT-H2B-FP-invFRT cassettes were assembled into the pJFRC-MUH backbone vector (Addgene, Inc.) by standard cloning method, so that the expression of each cassette is under the control of an upstream activation sequence (UAS) and an p10 poly-adenylation (p10pA) sequence (Pfeiffer et al., 2012). The final *LT-Bitbow* targeting plasmid was assembled by Gibson assembly (Gibson et al., 2009) into the same pJFRC-MUH backbone and was integrated into *Drosophila melanogaster* genome docking site VK00027 (BestGene, Inc.) using the ΦC31 integrase-mediated transgenesis systems (Bateman et al., 2006; Bischof et al., 2007; Groth et al., 2004; Markstein et al., 2008; Venken et al., 2006).

### *LT-Bitbow* sample preparation and imaging

*LT-Bitbow* transgenic flies, *LT-Bitbow*^20355-2M3^, was crossed with flies that carry heat-shock-inducible FLP^122^ and elav-Gal4, and raised at 23 °C for 2 days. To induce FP expression in the larval PNS, embryos were collected within consecutive 1-hour windows on day 3, and kept at 23 °C until heat shock at 2-3, 3-4, 4-5, 5-6, or 6-7 hr AEL. To induce FP expression in the larval CNS, 1^st^ instar larvae were heat-shocked on 1 day AEL. The heat shock was performed by heating the embryo-containing agar-plate on the surface of a 70 ℃ heat-block for 1.5 min, followed by a 30 min heat chamber incubation at 37 °C. The plate was then transferred back to the 23 °C incubator for the next 4 days.

Late 3^rd^ instar larvae were dissected to make fillet preparations in 1x hemolymph-like 3.1 saline buffer (Feng et al., 2004), immediately followed by fixation in 4% paraformaldehyde in 1x PBS for 20 min at room temperature (RT) with gentle shaking. Larval fillets were transferred to 1.5 ml Eppendorf tubes wrapped in tinfoil to prevent possible photobleaching by ambient lights. The samples were washed 3 times with 0.1% Triton X-100 in 1x PBS, 15 min each at RT, on a nutator, and then incubated in 5% normal donkey serum containing 0.1% Triton X-100 in 1x PBS for 30 min at RT. Alexa Fluor® 647 AffiniPure Goat Anti-Horseradish Peroxidase antibody (1:500, Jackson Immuno Research) was added to the samples for overnight incubation at 4 ℃ with gentle shaking. Samples were washed 3 times as described above. Fillets were placed on poly-L-lysine-coated coverslips and mounted with ProLong Diamond Antifade mountant (Invitrogen^TM^). Samples were left dry in the dark overnight at RT and imaged by confocal imaging on the following day or kept at 4 °C for up to a week until imaging.

*LT-Bitbow* samples were imaged with a Zeiss LSM780 laser scanning confocal microscope. PNS data were collected from abdominal segments 2 to 7 (A2-A7). Fiji with customized plugin scripts was used to process raw images for FP spectral unmixing (Schindelin et al., 2012). Individual FP signals assuming the shape of a nucleus in the soma were identified on a single z-plane. If a particular FP signal were observed in a neuron, it is recorded as an “1” in the color code; if that FP’s signal is not observed, it is then recorded as a “0”. Color code of each neuron was recorded manually to create a data sheet for each experimental condition, which were subsequently analyzed computationally and statistically.

### Quantification of lineage relatedness

The goal of our statistical methods was to systematically identify neurons that share a common lineage and those do not share a common lineage with any others. A key aspect of this test is the ability to score the lineage relatedness for a pair of two neurons based on the color codes recorded from the *LT-Bitbow* experiments. To do this, we implemented a modified version of the Cohen’s Kappa (κ) to quantify the degree of color-code agreement between two neurons, which is defined as:

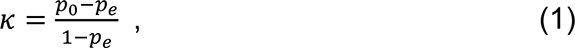

where *p*_0_ and *p*_*e*_ are the degree of color code agreement between the two neurons calculated by observation and by chance, respectively.

To calculate the κ value for each neuron pair, we first calculate the probability of a neuron *n* that expresses a particular color *c*, *P*_*n*,*c*_:

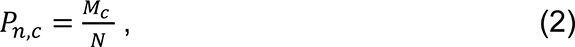

where, *M*_*c*_ is the total observation that a neuron expresses color *c*, and *N* is the total number of neurons. As shown in Figure 2D, the percentage of neuron expressed any fluorescent color was relatively low (∼6% maximum). We therefore hypothesize that most of the null-color (00000) neurons were the result of no Flp recombination at all, instead of due to the same FP being recombined even times. As a result, we estimated the κ in the cases that at least one of the neurons in a comparing pair needs to express at least one FP to avoid the false identification of null-color expressing neurons being lineage related. Based on this assumption, we can estimate the *p*_*e*_ value between neurons *i* and *j* (i.e., 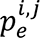), in which at least one of them expresses at least one FP. The 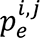 value can be calculated by summing the products of the *P*_*n*,*c*_ for both neurons for all the non-null colors (denoted from 1 to *C*):

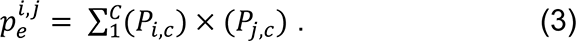

To calculate the *p*_0_ value of the corresponding neuron pair 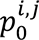, we again, only considered the cases in which at least one of the neurons expressed at least one FP:

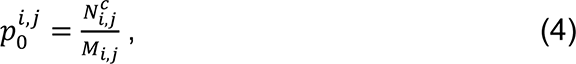

where, 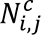 is the total observations that both neurons expressed a same non-null colors color and *M*_*i*,*j*_ is the total observations that either neuron expressed a non-null color.

Finally, we acquired the observed κ values for all the neuron pairs using their *p*_*e*_ and *p*_0_ values following equation (1). An analysis of this equation reveals that if any two neurons always have perfectly matched colors (i.e., *p*_0_ = 1), they will have a κ = 1. If two neurons’ colors are matched exactly as would be expected by chance (i.e., *p*_0_ = *p*_*e*_), then they will have a κ = 0. In this way, a higher κ value indicates a higher probability that the two neurons are labeled in the same color not by chance (i.e., lineage-related).

### Statistical analysis of lineage relatedness

We designed a statistical test to determine whether the experimentally observed κ value of a neuron pair is significantly higher than that by chance. To estimate the κ value distribution from labeling a particular neuron pair by random colors, we established a randomization process, starting with generating a raw color matrix where every row represents an individual PNS hemi-segment, and each column represents a neuron within that hemi-segment. The κ values were calculated for all possible combinations of neuron pairs within the data. Next, the raw color matrix was randomized by shuffling the colors each neuron expressing any non-00000 null-color. This ensures that the neurons receiving a color in the randomized data will have also had a color in the real data. We reason that this randomization process is more accurate than a total randomization process because each neuron pair remains the same total chance to match colors as in the real experiments, in which the ambiguous 00000 null-color are excluded from being assigning as a matched color (see above). After the color randomization, the κ values were calculated for all possible combinations of neuron pairs within the data. This randomization process was repeated 100,000 times for each neuron pair to generate a distribution of κ values that were compared to the κ value obtained from the experimental observation. The number of times that the random κ value was greater than or equal to the observed κ value was counted and then divided by 100,000 to generate a p-value to estimate whether the experimentally observed κ value is significantly larger than those calculated by random color assignment. Finally, the p-values was adjusted for multiple hypothesis testing by calculating a false discovery rate (FDR) using the Python statsmodels package (https://www.statsmodels.org/dev/generated/statsmodels.stats.multitest.multipletests.html). The Benjamini/Hochberg (non-negative) method was used for FDR evaluation (Benjamini and Hochberg, 1995).

It is important to run a separate test for each neuron pair because certain neurons expressed a color more often than others. This allowed us to normalize the high expressing neurons in a way that they did not just have a better κ value by chance.

The implementation of our lineage relatedness statistical model is available at https://github.com/MikeVeling/Process-Brainbow-Data.

The complete lineages map of PNS neurons was generated by the Cytoscape Software package (version 3.7.1., https://cytoscape.org/index.html).

### *dar1 cis*-reporter constructs and transgenic fly generation

Genomic DNA (gDNA) of the *white^1118^* strain of *Drosophila melanogaster* was extracted by standard method, and used as template for amplifying the 500 bp and 100 bp *cis*-regulatory elements with restriction sites (BamHI and EcoRI) added to the 5’ and 3’ end, respectively. The amplified DNA fragments were each cloned into the gateway entry vector (pBP-ENTR/D-TOPO) via restriction digestion and T4 ligation reaction. Subsequently the Invitrogen^TM^ Gateway® LR clonase® II system was used to integrate the *cis*-element entry vectors into the destine backbone vector pBPGUw, which carries a minimal promoter to allow expressing Gal4 in the presence of active *cis*-regulatory element(s) (Pfeiffer et al., 2008; Pfeiffer et al., 2010; Swanson et al., 2008). Control transgene was made in the same backbone vector without adding any *cis*-regulatory element into the gateway entry vector. Purified plasmids were integrated into *Drosophila* genome at the attP40 docking site using the ΦC31 integrase-mediated transgenesis systems (Bateman et al., 2006; Bischof et al., 2007; Groth et al., 2004; Markstein et al., 2008; Venken et al., 2006).

### Sample preparation and imaging for *dar1* expression reporters

*dar1 cis*-element-Gal4 transgenic flies were mated with UAS-GFP reporter flies (UAS-GFP^VK33^ or UAS-lacZ.nls::GFP) at 25 °C. The late 3^rd^ instar larvae from the cross were dissected to make fillet preparations, fixed, washed, and blocked for immunostaining as described above for *LT-Bitbow* samples. Primary antibodies chicken anti-GFP (Aves, 1:1000 dilution) and rat anti-Elav (Developmental Studies Hybridoma Bank, 1:100 dilution) were added to larval fillets for overnight incubation at 4 °C. Samples were then washed 3 times and then incubated with secondary antibodies for overnight incubation at 4 °C. The secondary antibodies used were Alexa Fluor® 488 AffiniPure Donkey Anti-Chicken IgY (IgG) (H+L), Rhodamine Red™-X (RRX) AffiniPure Donkey Anti-Rat IgG (H+L), and Alexa Fluor® 647 AffiniPure Goat Anti-Horseradish Peroxidase (Jackson Immuno Research, 1:500 dilution). After being washed 3 times, the samples were placed on poly-l-lysine coated coverslips for ethanol dehydration at 30%, 50%, 70%, 95%, 100% (two times) ethanol for 5 minute each. After that, the samples were treated with 100% xylene twice for 10 minute each. The fillet samples were mounted with DPX mounting media (Electron Microscopy Sciences) and left dry in the dark for at least overnight at RT. Confocal imaging was performed with a Leica TCS SP5 confocal microscope. PNS neurons in segments A2 to A7 were imaged.

