## Supplemental Information for "Identification of neuronal lineages in the *Drosophila* peripheral nervous system with a novel multi-spectral lineage tracing system"

### Inventory of Supplemental Information

#### Supplemental Data

Figure S1.
related to Figure 3.

Figure S2.
related to Figure 3.

Figure S3.
related to Figure 4.

Figure S4.
related to Figure 5.

#### Supplemental Figure Legends

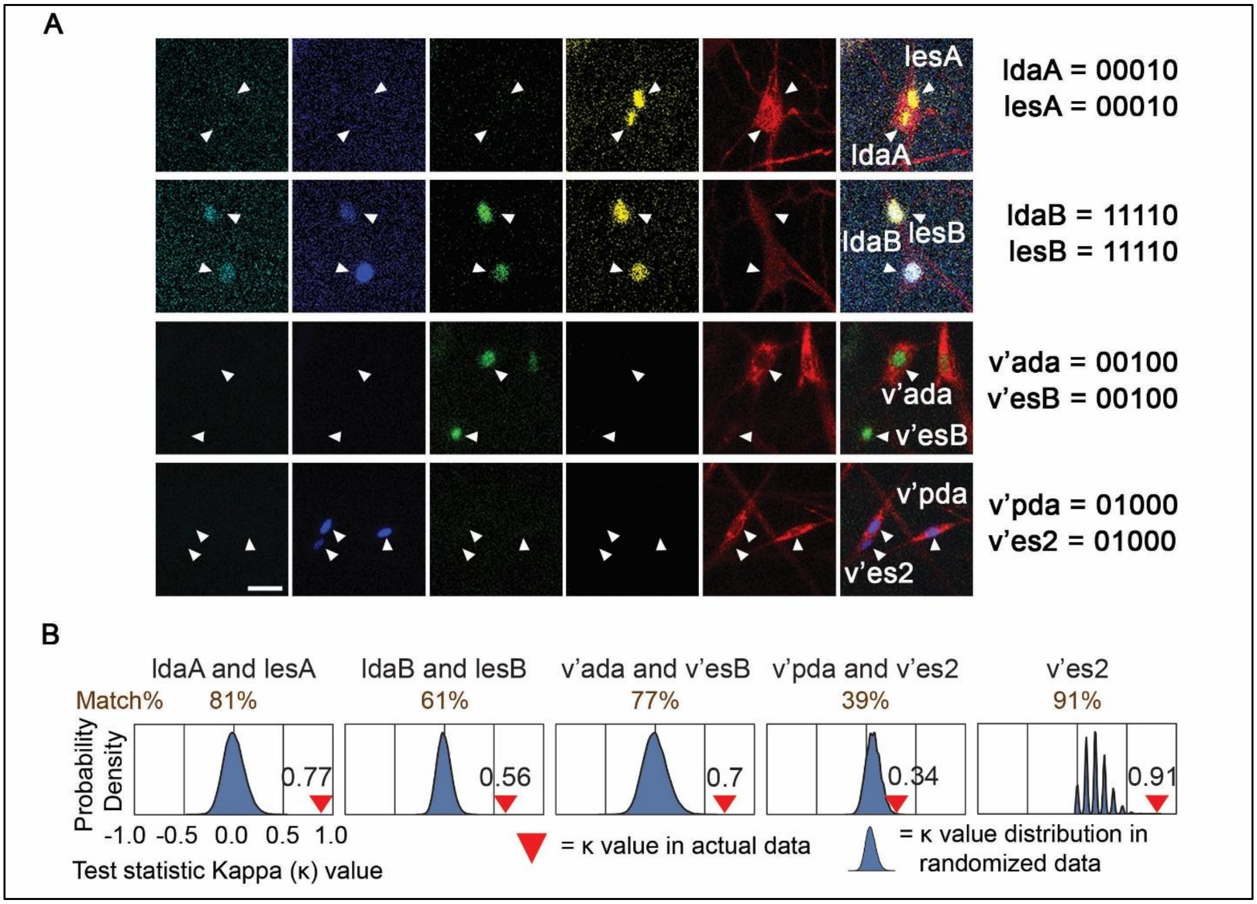


**Figure S1 (related to Figure 3). Lineage tracing by *LT-Bitbow* confirms some previously reported lineages.**
(**A**) Representative fluorescent images for the previously reported PNS neuron lineages ldaA-lesA, ldaB-lesB, v'ada-v'esB, and v'pda-v'es2 (Brewster and Bodmer, 1995). The color codes of the fluorescent proteins expressed in the neurons are shown on the right. Scale bar: 10 μm.
(**B**) The neuron pairs in these lineages have significantly higher positive κ values calculated from the experimental observations than those calculated from the randomized model in the relatedness tests.

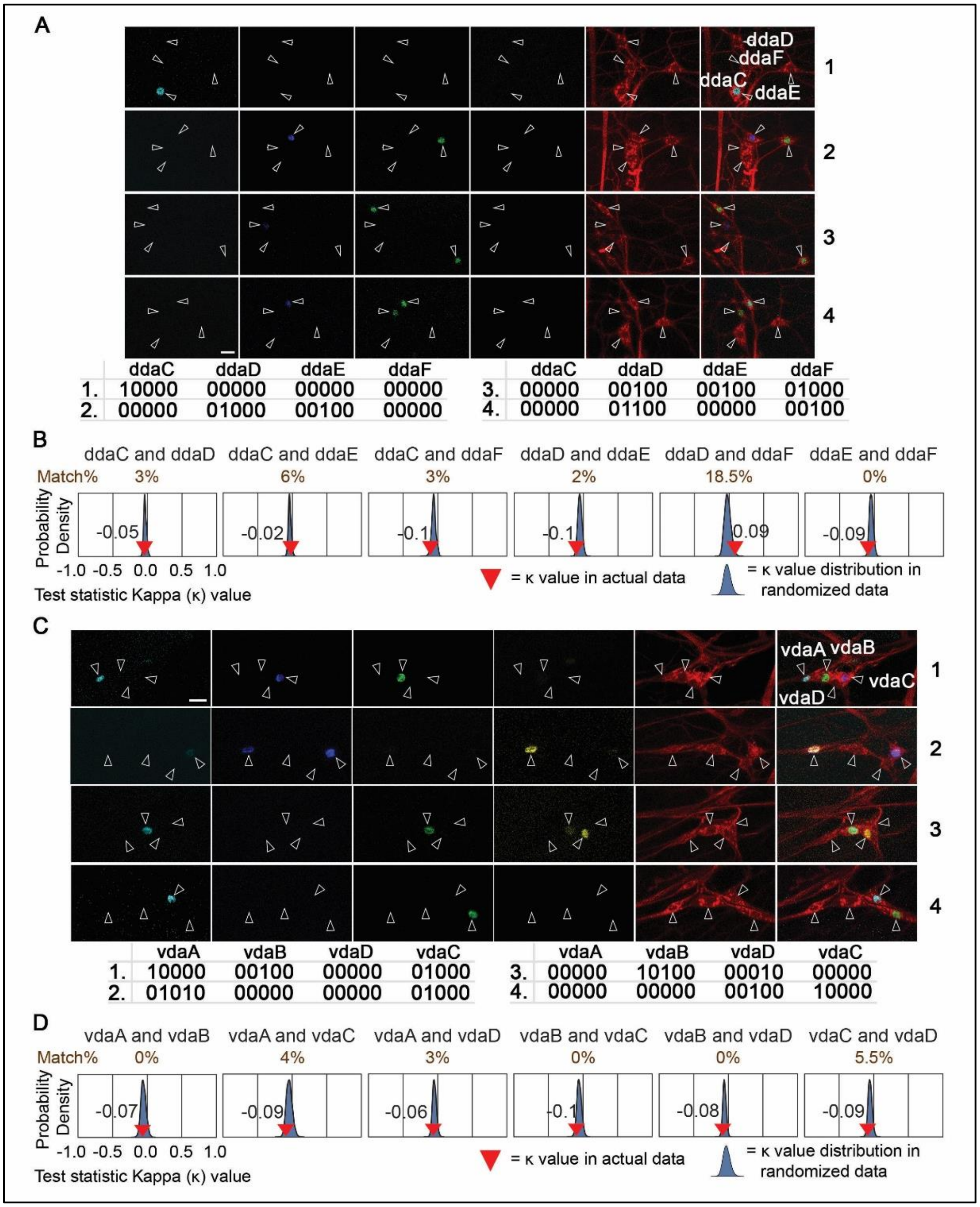


**Figure S2 (related to Figure 3). Lineage tracing by *LT-Bitbow* rejects some previously reported lineages.**
(**A**) Representative fluorescent images for the previously reported lineage-related neurons in the dmd cluster (Brewster and Bodmer, 1995). The color codes of the fluorescent proteins expressed in the neurons are shown at the bottom. Scale bar: 10 μm.
(**B**) The κ values of the neuron pairs in the dmd cluster are not significantly different from those calculated with the randomized model in the relatedness tests.
(**C**) Representative fluorescent images for the previously reported lineage-related neurons in the vmd cluster (Brewster and Bodmer, 1995).
(**D**) The κ values of the neuron pairs in the vmd cluster are not significantly different from those calculated with the randomized model.

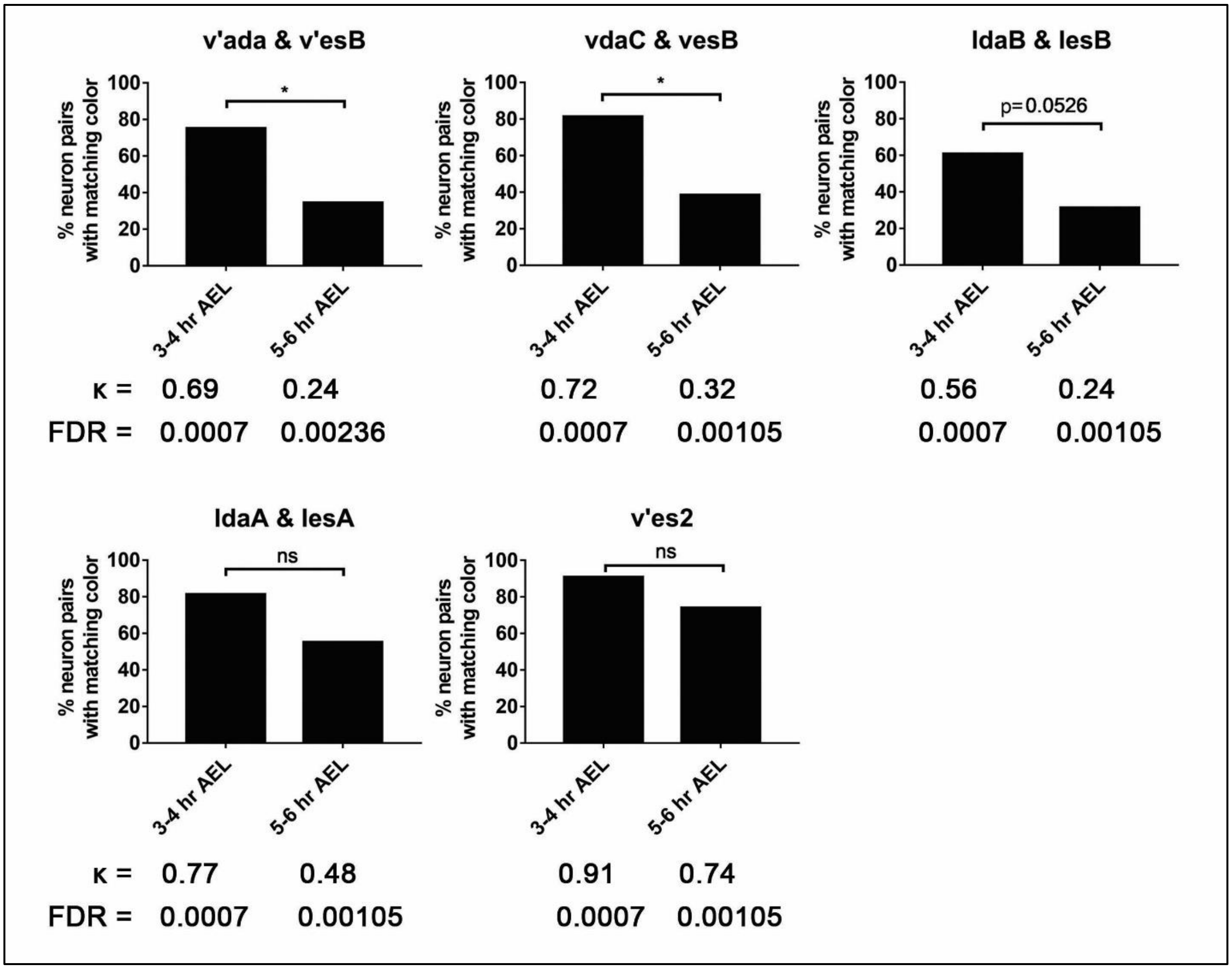


**Figure S3 (related to Figure 4).** ***LT-Bitbow*** **provides an efficient way to compare the birth-time of neurons in different lineages.**
The percentage of neuron pairs with matching colors decreases significantly for several lineages (e.g., v'ada-v'esB, vdaC-vesB, ldaB-lesB) but remains high for some lineages (e.g., the ldaA-lesA lineage, and two neurons in the v'es2 group), suggesting the former are closer to finishing their lineage development than the latter. The percentage of neuron pairs with matching colors is quantified by dividing the number of total neuron pairs that have the same colors to the number of total neuron pairs that are labeled by any color. All neuron pairs analyzed in this experiment are lineage-related, as determined by the relatedness test. *: $\alpha < 0.05$, Fisher's exact test.

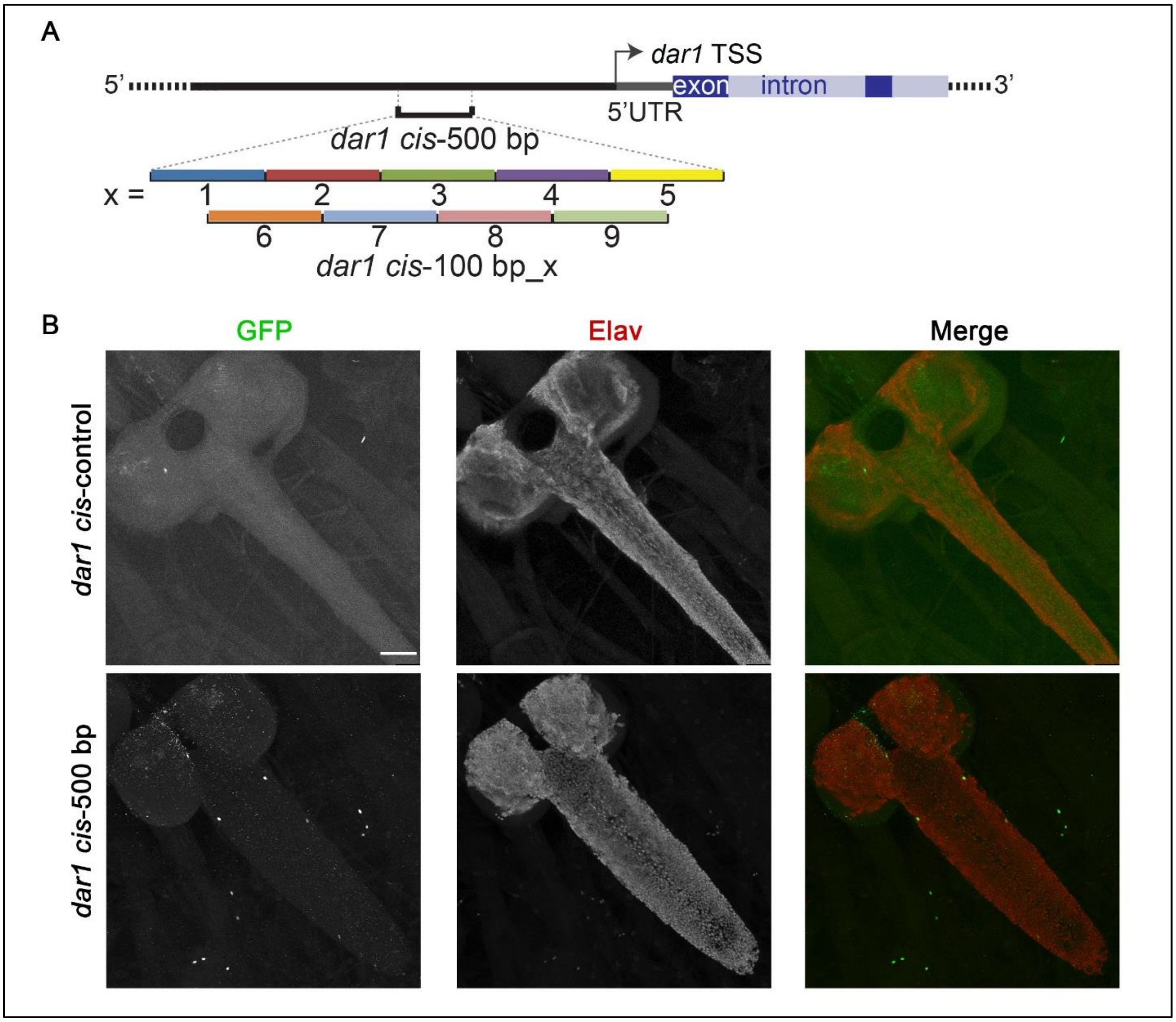


**Figure S4 (related to Figure 5). Identification of the *cis*-regulatory elements of *dar1* expression by using GFP reporters.**
(**A**) Schematic drawing that shows the genomic locations of the 500-bp fragment that largely recapitulates *dar1* expression and the 100-bp fragments used to further identify the *cis*-regulatory elements. TSS: transcription start site.
(**B**) Both the control driver and the 500-bp *dar1* fragment drive the expression of the GFP-reporter only in a few neurons in the entire CNS, which contains only non-multipolar neurons. Scale bars: 50 μm.
